# Autophagy adaptors mediate Parkin-dependent mitophagy by forming sheet-like liquid condensates

**DOI:** 10.1101/2023.09.11.557117

**Authors:** Zi Yang, Saori R. Yoshii, Yuji Sakai, Haruka Chino, Roland L. Knorr, Noboru Mizushima

## Abstract

During PINK1 and Parkin-mediated mitophagy, autophagy adaptors are recruited to depolarized mitochondria to promote the selective degradation of mitochondria. Autophagy adaptors such as OPTN and NDP52 bridge mitochondria and autophagosomal membranes by binding to ubiquitinated mitochondrial proteins and autophagosomal ATG8 family proteins. Here, we demonstrate that OPTN and NDP52 form sheet-like phase-separated condensates with liquid-like properties on the surface of ubiquitinated mitochondria. The dynamic and liquid-like feature of OPTN condensates is important for mitophagy activity because reducing the liquidity of OPTN–ubiquitin condensates suppresses the recruitment of ATG9 vesicles and impairs mitophagy. Based on these results, we propose a dynamic liquid-like model of autophagy adaptors, in contrast to a stoichiometric model, to explain their interactions between autophagic membranes (i.e., ATG9 vesicles and isolation membranes) and mitochondrial membranes during Parkin-mediated mitophagy. This model underscores the importance of liquid–liquid phase separation in facilitating membrane– membrane contacts, likely through the generation of capillary forces.

## Introduction

Macroautophagy (hereafter referred to as autophagy) is an intracellular degradation process mediated by a double-membraned organelle termed the autophagosome (Nakatogawa, 2020; Mizushima & Levine, 2020). Cytosolic components such as proteins and organelles are segregated into autophagosomes, which subsequently fuse with lysosomes for degradation. Autophagy can be either non-selective or selective (Lamark & Johansen, 2021; Vargas *et al*, 2023). Mitophagy, which is a selective type of autophagy, specifically degrades mitochondria and can be induced by mitochondrial damage or cellular stresses (Onishi *et al*, 2021; Ganley & Simonsen, 2022). For example, after losing the mitochondrial membrane potential, numerous mitochondrial outer-membrane proteins are ubiquitinated by the PINK1-Parkin system, initiating the recruitment of autophagy adaptors, including optineurin (OPTN), calcium binding and coiled-coil domain 2 (CALCOCO2/NDP52), sequestosome 1 (SQSTM1/p62), Next to BRCA1 gene 1 protein (NBR1), and Tax1 binding protein 1 (TAX1BP1) (Lamark & Johansen, 2021; Vargas *et al*, 2023). These autophagy adaptors bridge the mitochondrial and autophagosomal membranes by interacting with mitochondrial ubiquitinated proteins via a ubiquitin-binding domain (UBD) and with autophagosomal ATG8 family proteins (i.e., LC3 and GABARAP family proteins in mammals) via the LC3-interacting region (LIR). In addition to their role in bridging mitochondria and autophagosomes, OTPN and NDP52 also play crucial roles in inducing mitophagy by recruiting upstream autophagy factors (Lazarou *et al*, 2015); OPTN recruits ATG9 vesicles, whereas NDP52 recruits the ULK1 complex through interaction with FIP200 (Yamano *et al*, 2020; Vargas *et al*, 2023).

Liquid–liquid phase separation (LLPS) has been described in various biological processes (Musacchio, 2022; Hirose *et al*, 2023). LLPS involves the formation of liquid-like condensates through dynamic, multivalent interactions between various molecules that often possess intrinsically disordered regions. Emerging evidence suggests connections between autophagy and LLPS at multiple steps, including autophagosome formation and degradation (Noda *et al*, 2020; Ma *et al*, 2023). Notably, autophagy adaptors, such as p62, undergo LLPS through multivalent interaction with poly-ubiquitins and form cytosolic condensates (Zaffagnini *et al*, 2018; Sun *et al*, 2018). These condensates can be degraded by autophagy, which is termed “fluidophagy” (Agudo-Canalejo *et al*, 2021). Furthermore, recent studies showed that p62 undergoes LLPS on mitochondria and lysosomes that are subjected to selective autophagy (Peng *et al*, 2021; Gallagher & Holzbaur, 2023).

Generally, phase-separated condensates can deform when they contact a rigid surface in order to minimize the overall energy of the system, a phenomenon known as “wetting” (Kusumaatmaja *et al*, 2021b; Gouveia *et al*, 2022). The wetting behavior of intracellular condensates is determined by the interfacial tensions between the condensate–membrane, condensate–cytosol, and cytosol-membrane interfaces. Depending on the relative strengths of these interfacial tensions, the condensates exhibit partial wetting, complete wetting, or de-wetting. In contrast, elastic surfaces, including membranes to which phase-separated condensates adhere, also deform (Kusumaatmaja *et al*, 2021a, 2021b; Gouveia *et al*, 2022). This indeed happens during fluidophagy; autophagosomal membranes bend along the surface of p62 condensates (Agudo-Canalejo *et al*, 2021). Deformation of a membrane by a phase-separated condensate is determined by the energy balance between membrane deformation and the surface tension of the condensate, which is known as elastocapillarity (Style *et al*, 2017; Kusumaatmaja *et al*, 2021b).

By forming capillary bridges, wetting droplets can also induce adhesion of surfaces (Wexler *et al*, 2014). Whether LLPS and capillary bridges also contribute to the adhesion of cellular membranes is not presently known. Therefore, we hypothesized that phase-separated condensates of autophagy adaptors can wet both mitochondria and autophagic membranes, including ATG9 vesicles and isolation membranes (also called phagophores), thereby promoting their contact during Parkin-mediated mitophagy. In this study, we employ live-cell imaging and a mathematical model to provide evidence that the autophagy adaptors form phase-separated condensates on the surface of the mitochondrial membrane, resulting in sheet-like condensates that cover the surface of ubiquitinated mitochondria upon mitophagy induction. These condensates accumulate between mitochondria or between mitochondria and autophagic membranes, exhibiting a dynamic nature with a liquid-like property. Furthermore, we demonstrate the essential role of these dynamic condensates in the recruitment of ATG9 vesicles and the initiation of mitophagy. Based on these results, we propose a dynamic liquid-like model, rather than a stoichiometric model, to describe the roles of autophagy adaptors in Parkin-mediated mitophagy.

## Results

### Autophagy adaptors show distinct distributions during Parkin-mediated mitophagy

To better understand the role of each autophagy adaptor, we analyzed their localization during Parkin-mediated mitophagy. When mitophagy was induced by treatment with the mitochondrial uncoupler carbonyl cyanide *m*-chlorophenylhydrazone (CCCP) in HeLa cells expressing exogenous Parkin, the autophagy adaptors p62, OPTN, NBR1, NDP52, and TAX1BP1 translocated to mitochondria as previously observed (Fig. 1A and B) (Geisler *et al*, 2010; Wong & Holzbaur, 2014; Heo *et al*, 2015; Moore & Holzbaur, 2016; Gallagher & Holzbaur, 2023). Ubiquitin and all of these autophagy adaptors were distributed evenly on the surface of separate mitochondria (Mt; Fig. 1A). In contrast, these adaptors exhibited distinct localization patterns during the formation of isolation membranes on mitochondria (Mt–IM; Fig. 1B and C). The signals of OPTN and NDP52 were enriched in areas where LC3B signals colocalized, while they were mostly absent from the LC3B-negative side of the same mitochondria (Fig. 1B and C). Endogenous OPTN and NDP52 also showed clear enrichment on the LC3B-positive areas in comparison to the LC3B-negative side, confirming that this localization is not caused by the overexpression of adaptors (Fig. 1D). However, this inhomogeneous distribution pattern was not observed for ubiquitin, p62, NBR1, and TAX1BP1 (Fig. 1B and C). Unlike OPTN and NDP52, p62 showed enrichment between clustered mitochondria (Mt-Mt), which is consistent with previous reports (Fig. EV1A and B) (Wong & Holzbaur, 2014). These accumulation patterns cannot be explained by the stoichiometric interaction of the autophagy adaptors with ubiquitinated proteins that are evenly distributed on mitochondria.

**Figure 1.**
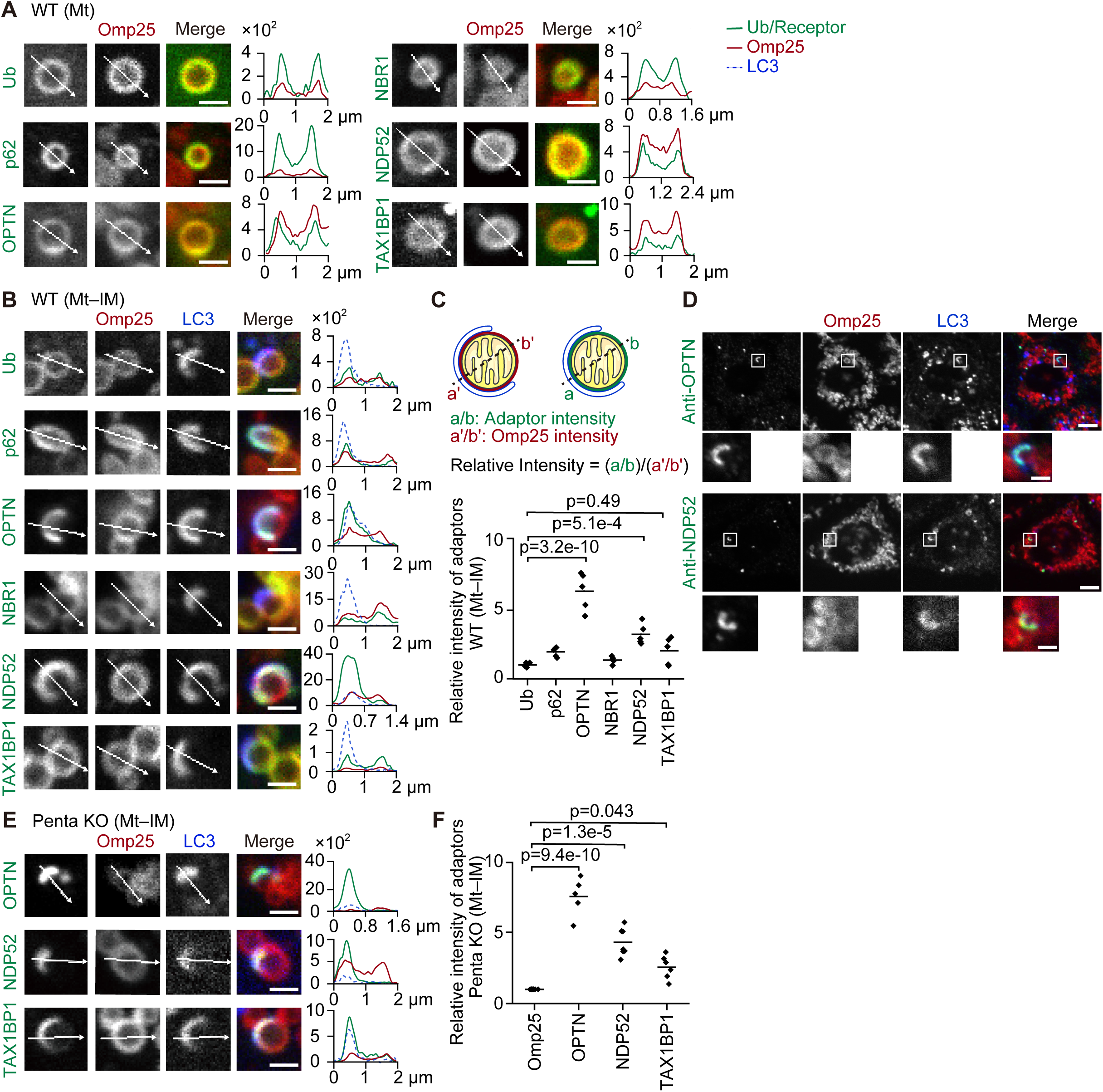
Autophagy adaptors show distinct distributions during Parkin-mediated mitophagy. (A, B, and E) Representative images (left) and spline graphs of the intensity profiles along the indicated lines in the respective images (right) of wild-type HeLa cells (A and B) or HeLa cells lacking all five autophagy adaptors (penta KO cells) (E) expressing one of the GFP-tagged autophagy adaptors or ubiquitin, mRuby–Omp25, and HaloTag7–LC3B at 45 min after 20 µM CCCP treatment. Separate mitochondria without contacting other mitochondria or isolation membranes (A; Mt) or mitochondria with an isolation membrane (B and E; Mt–IM) are shown. The *y*-axis in each of the graphs indicates the fluorescence intensity. Scale bars indicate 1 μm. (C and F) The relative intensity of GFP-tagged adaptors or ubiquitin (a and b) in LC3B-positive areas compared with LC3B-negative areas in panels B and E. GFP signals were normalized to the mRuby–Omp25 signals (a′ and b′) using the formula Relative Intensity = (a/b)/(a′/b′), where a and a′ represent the intensities in the LC3B-positive area, and b and b′ represent the intensities in the LC3B-negative area. Solid horizontal bars indicate the means, and dots indicate the data from five structures. Differences were statistically analyzed by one-way analysis of variance with Dunnett’s post-hoc test. (D) Immunostaining of endogenous OPTN or NDP52 in wild-type HeLa cells expressing mRuby–Omp25 and HaloTag7–LC3B at 45 min after 20 µM CCCP treatment. Scale bars indicate 4 µm and 1 µm (magnified images).

The localization of autophagy adaptors may be affected by their interaction with other adaptors (Turco *et al*, 2021; Gubas & Dikic, 2022). To examine the localization of each adaptor on its own, we used HeLa cells lacking all five autophagy adaptors (penta KO cells) to exclude the effect of other adaptors (Lazarou *et al*, 2015). In penta KO cells, OPTN and NDP52 exhibited significant enrichment in LC3B-positive areas, similar to that observed in wild-type cells, suggesting that this accumulation occurred independently of heterologous interaction with other adaptors (Fig. 1E and F). Mitophagy was not restored by exogenous expression of p62 or NBR1 in penta KO cells, and thus colocalization with LC3 could not be tested for these two adaptors (Lazarou *et al*, 2015). The accumulation of p62 between clustered mitochondria was also observed in penta KO cells (Fig. EV1C and D). Given the uniform distribution of ubiquitinated proteins on mitochondrial surfaces, these data suggest that autophagy adaptors exhibit non-stoichiometric enrichment between membranes. Hereafter, we used penta KO cells to study each autophagy adapter individually

### OPTN and NDP52 show a dynamic exchange between the mitochondrial surface and cytosol

We hypothesized that the non-stoichiometric distribution of the autophagy adaptors on ubiquitinated mitochondria could be explained by LLPS. Typical condensates generated by LLPS exhibit rapid exchange of their components, which is often demonstrated by fluorescence recovery after photobleaching (FRAP) analysis (Taylor *et al*, 2019; McSwiggen *et al*, 2019). We first conducted FRAP experiments for separate (i.e., unclustered) mitochondria in CCCP-treated cells. As expected, ubiquitin showed virtually no recovery because it was conjugated to mitochondrial membrane proteins (Fig. 2A and B). Additionally, p62 recovered only slightly, suggesting a minute exchange between the mitochondrial surface and cytosol (Fig. 2A and B). In contrast, OPTN and NDP52 showed a rapid recovery, indicating a dynamic exchange of these adaptors between the mitochondrial surface and the cytosol (Fig. 2A and B).

**Figure 2.**
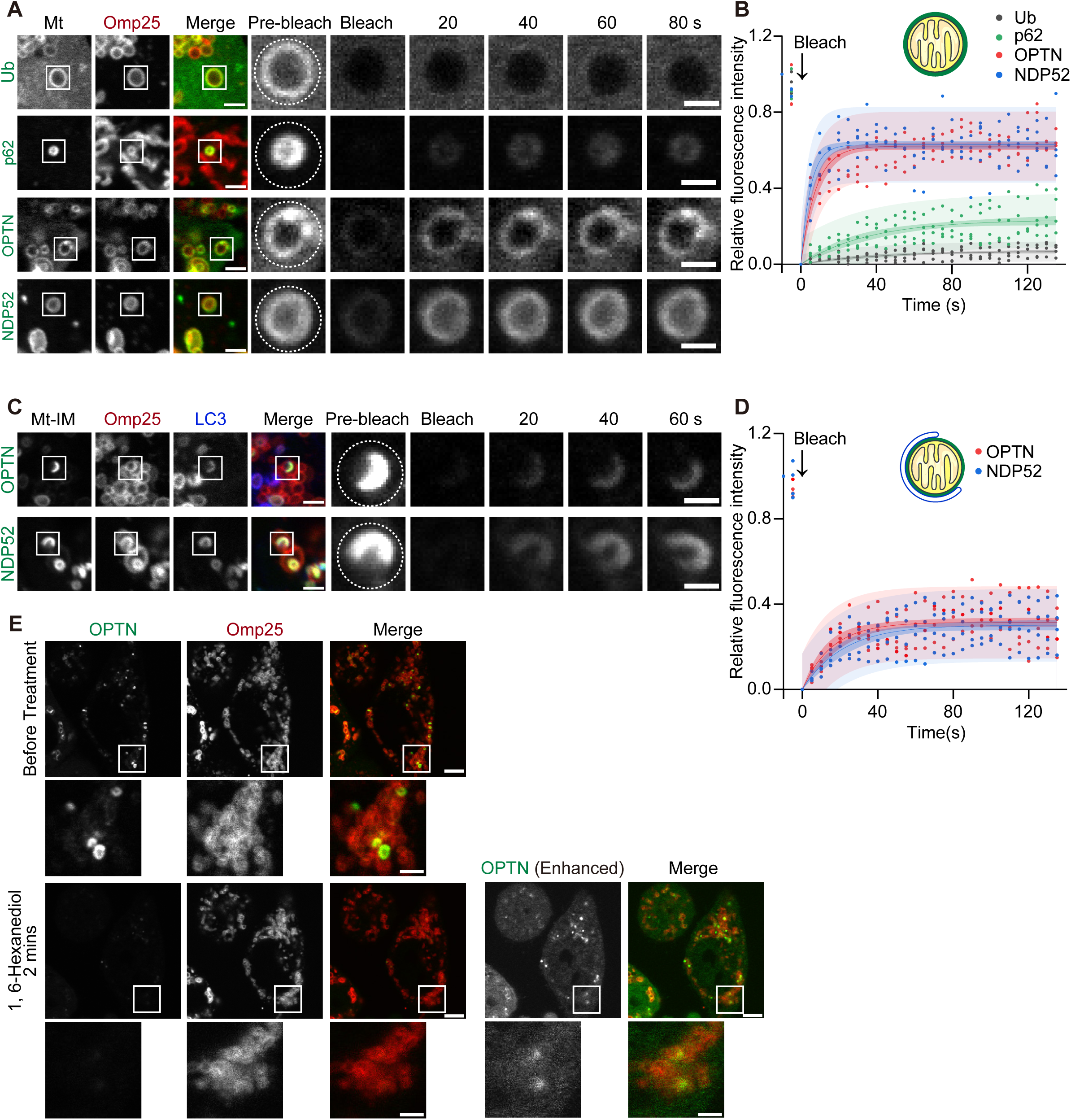
OPTN and NDP52 show a dynamic exchange between the mitochondrial surface and cytosol. (A–D) Representative images (A and C) and quantification (B and D) of GFP fluorescence recovery after photobleaching (FRAP) on separate mitochondria (Mt) (A and B) or on mitochondria with an isolation membrane (Mt–IM) (C and D) in HeLa cells lacking all five autophagy adaptors (penta KO cells) expressing GFP-tagged adaptors or ubiquitin at 45 min after 20 µM CCCP treatment. The photobleached areas are indicated by dotted lines. The magnified panels display time-lapse images of the photobleached areas. Scale bars indicate 2 μm and 1 μm (magnified images). Data were collected from four structures and were fitted to the equation *y* = *a*\*(1 − exp(−*b*\**x*)). The dark shaded areas represent the 95% confidence intervals, and the light shaded areas represent the 95% prediction intervals. (E) Penta KO cells expressing GFP–OPTN at 45 min after 20 µM CCCP and 20 μM wortmannin treatment. Wortmannin was added to inhibit autophagosome formation so that OPTN would not be sequestered in a closed compartment. Images of cells before (upper panels) and 2 min after (lower panels) the addition of 10% 1,6-hexanediol are displayed. Scale bars indicate 5 μm and 2 μm (magnified images).

Next, we examined the dynamics of adaptors between mitochondria and isolation membranes. OPTN and NDP52 showed partial recovery (Fig. 2C and D). Although this result indicates some exchange of these adaptors between membranes, the incomplete recovery suggests the existence of a gel-like or immobile fraction. Treatment with 1,6-hexanediol, which can be used to dissolve condensates, dispersed OPTN from mitochondria without isolation membranes, suggesting that OPTN accumulation is supported by weak interactions (Fig. 2E). These results indicate that OPTN and NDP52 on ubiquitinated mitochondria are mobile and can quickly exchange between the mitochondrial surface and the cytosol, supporting the hypothesis that they form phase-separated condensates on damaged mitochondria.

### Mathematical models of condensate formation and localization of autophagy adaptors

To simulate the behavior of autophagy adaptors in the context of phase-separated condensates, we developed mathematical models (see Materials and Methods for details). First, we considered the case of two mitochondria approaching each other (Fig. 3A). In Parkin-mediated mitophagy, various proteins in the mitochondrial outer membrane are ubiquitinated. We modeled the mitochondrial outer membrane as a circle with a diameter of D_Mt_ = 600 nm. On these mitochondria, we assumed a ring-shaped ubiquitin layer with a thickness w_*Ub*_ = 20 nm and area-fraction (hereafter referred to as concentration) *ϕ*_*Ub*_ = 0.1 (Milo & Phillips, 2015; Berry *et al*, 2018) (Fig. 3A). The closest distance between the two mitochondrial outer membranes was set to *d*_Mt_ (Fig. 3A). Here, we considered the dynamics of concentration changes of the adapter proteins and solvent components. We assumed that mitochondria were very large compared with proteins and did not move on a similarly short time scale, and that ubiquitin was tightly bound to mitochondrial membranes and immobile, as suggested by the FRAP experiment (Fig. 2B). The adaptor proteins were assumed to diffuse freely in solution and to bind to ubiquitin and themselves (self-interactions) with strengths *χ*_*Ub*_ and *χ*_*self*_, respectively.

**Figure 3.**
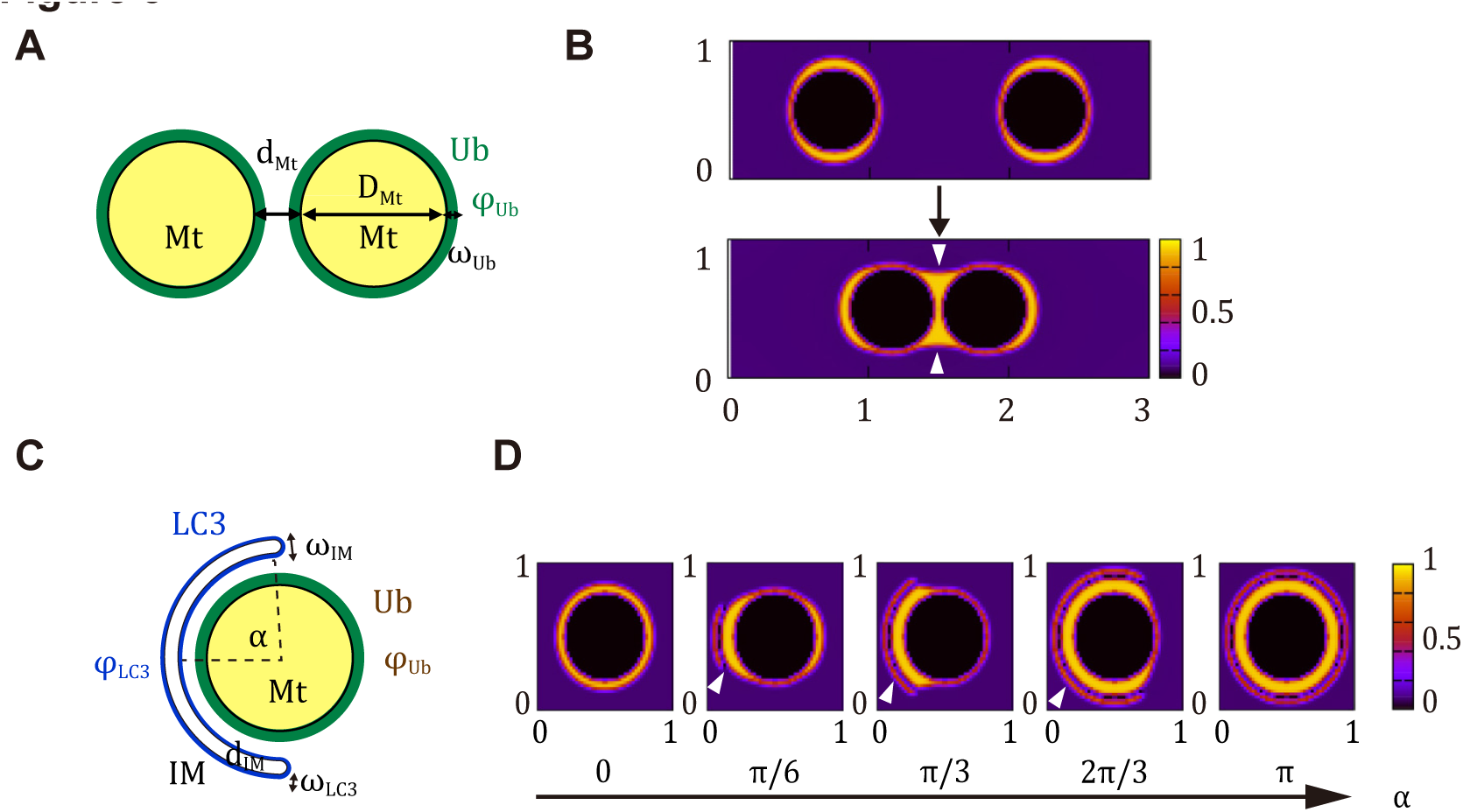
Mathematical models of condensate formation and localization of autophagy adaptors. (A) Schematic diagram of two separated mitochondria. (B) Sheet-like condensates of autophagy adaptors on mitochondria deform when the inter-mitochondrial distance *d_Mt_*was changed. The strength of adaptor self-interaction (*χ_self_*), adaptor–ubiquitin interaction (*χ_Ub_*), mitochondrial area exclusion (*χ_Mt_*) and surface tension (σ) were set to *χ_self_ =* 3*k_B_T*, *χ_Ub_* = 12 *k_B_T*, *χ_Mt_ =* 5*k_B_T*, and *σ* = *k_B_T*, respectively. The color indicates the adaptor concentration (*ϕ_ad_*). Arrowheads indicate autophagy adaptors filling the cleft between the two mitochondria. (C) Schematic diagram of a single mitochondrion with an isolation membrane. (D) Sheet-like condensates of autophagy adaptors on mitochondria deformed when the isolation membrane (arrowheads) appeared and the angle α surrounding the mitochondria changed. The strength of adaptor self-interaction (*χ_self_*), adaptor–ubiquitin interaction (*χ_Ub_*), adaptor–LC3 interaction (*χ_LC3_*), mitochondrial area exclusion (*χ_Mt_*), isolation membrane area exclusion (*χ_IM_*), and surface tension (σ) were set to *χ_self_ = 3k_B_T*, *χ_Ub_* = *χ_LC3_* = 12*k_B_T*, *χ*_Mt_ = *χ*_*IM*_ = 5*k_B_T*, and σ = *k_B_T*, respectively. The color indicates the adaptor concentration (*ϕ*_*ad*_).

Using the mathematical model, we assessed how the condensates deformed when the distance between two mitochondria *d*_Mt_ was changed, where *χ*_*Ub*_ = 12*k_B_T* and *χ*_*self*_ = 3*k_B_T*, with the Boltzmann constant *k_B_* and temperature *T*. When two mitochondria were far apart (*d*_Mt_ = 900 nm), the model predicted that adaptors cover the entire mitochondrial surface (Fig. 3B). When the mitochondria were close enough to come into contact (*d*_Mt_ = 60 nm), the adaptors moved to the area between the mitochondria and filled the cleft between them (arrowheads, Fig. 3B).

Next, we considered the case in which a growing isolation membrane surrounded and enclosed a single mitochondrion (Fig. 3C and D). A mitochondrion with ubiquitin was modeled as mentioned above, and the isolation membrane was modeled as a region of thickness w_*IM*_ = 50 nm bound by two semicircles with the same center as the mitochondrion. The closest distance between the isolation membrane and the mitochondrial outer membrane was *d*_*IM*_ = 100 nm, and the isolation membrane bent at an angle α to surround the mitochondrion. The surface of the isolation membrane was assumed to be covered by an LC3 region with a thickness *w*_*LC3*_ = 20 nm and concentration *ϕ*_*LC3*_ = 0.1. The adaptors interacted with both LC3 and ubiquitin with strengths *χ*_*LC3*_ and *χ*_*Ub*_, respectively.

Using this model, we determined the distribution of the adaptors when the isolation membrane appeared and elongated (changing the angle α surrounding the mitochondrion). The interaction strengths were set to *χ*_*Ub*_ = *χ*_*LC3*_ = 12*k_B_T* and *χ*_*self*_ = 3*k_B_T*. In the absence of the isolation membrane, the adaptors wet the entire mitochondrial surface uniformly (Fig. 3D). However, with the appearance of the isolation membrane, the adaptors remobilized to the area between the isolation membrane and mitochondrion, and this adaptor-enriched region elongated together with the isolation membrane. These modeling results suggest that the distribution of the adaptors on a mitochondrion can change upon contact with another mitochondrion or an isolation membrane.

### OPTN condensates redistribute upon membrane contact

Our mathematical models predict that the OPTN distribution changes when mitochondria cluster or are engulfed by isolation membranes. We then validated these predictions by using live-cell imaging. When two small mitochondria with uniform OPTN distributions on their surface approached each other, OPTN redistributed and formed a smooth surface that covered the two adjacent mitochondria (Fig. 4A–D, Movie EV1). Notably, strong OPTN enrichment was observed in the cleft between the two mitochondria (arrowheads, Fig. 4A and C), showing a pattern distinct from that of the outer membrane protein Omp25 (OPTN signal peaks localized outside of Omp25 signal peaks) (Fig. 4B and D). This phenomenon was apparent when the size of both (Fig. 4A) or one (Fig. 4C, Movie EV1) of the two mitochondria was less than 1 µm. Although OPTN formed sheet-like rather than spherical condensates, this phenomenon appeared to be similar to condensate coalescence, which is one of the hallmarks of liquid-like condensates (Hyman *et al*, 2014).

**Figure 4.**
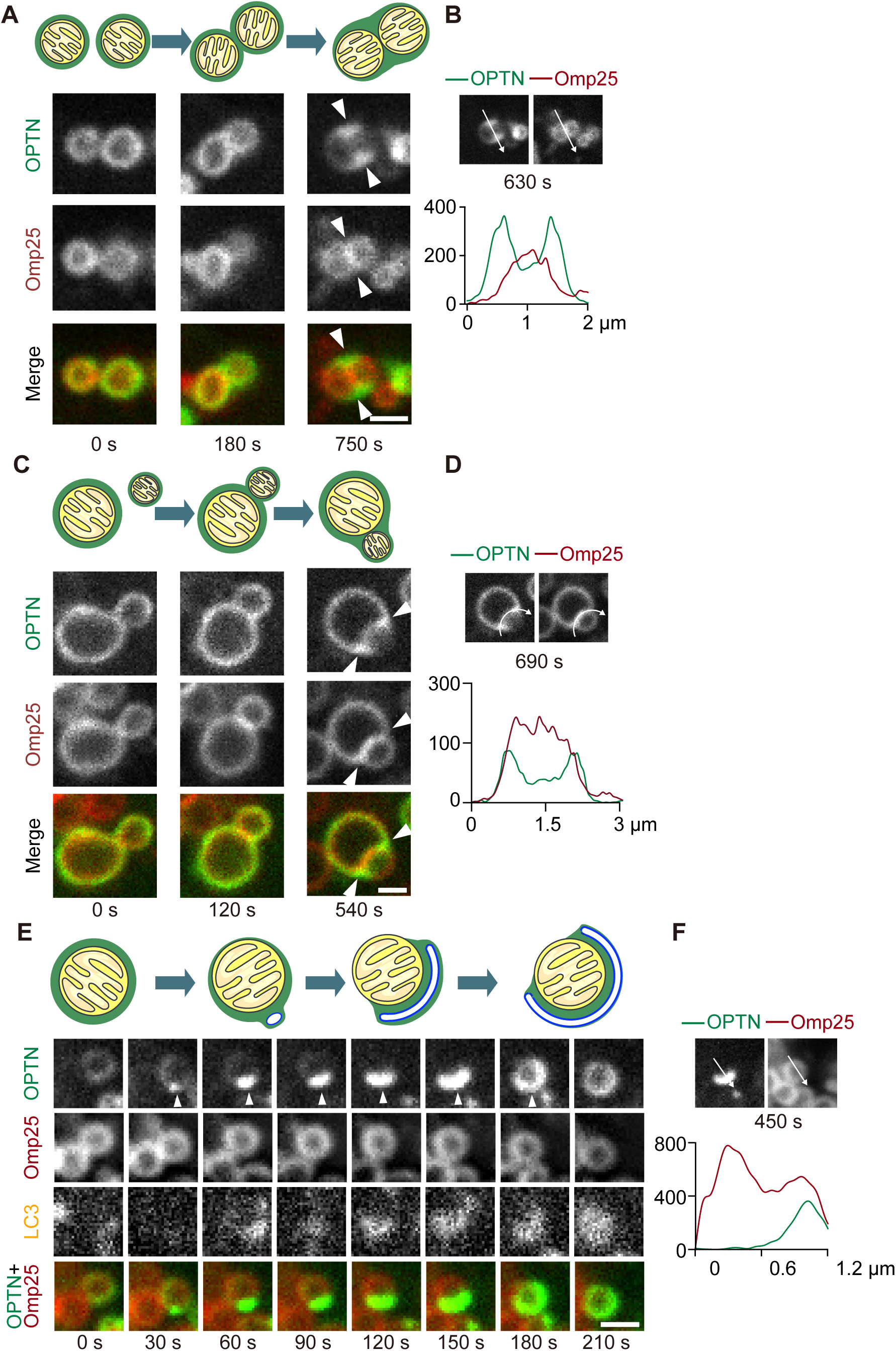
OPTN condensates redistribute upon contact with membranes. (A–D) Time-lapse images (A and C) and spline graphs (B and D) of two separate mitochondria approaching and contacting each other. Events of the attachment of two mitochondria with comparable sizes (A and B) and with differing sizes (C and D) were observed. The line graphs represent the intensity profiles along the indicated lines in the respective images shown (B and D). The arrowheads indicate redistributed OPTN signals filling the cleft between two adjacent mitochondria. Scale bars indicate 1 μm. See also Movie EV1 for the time-lapse video corresponding to (C). (E and F) Time-lapse images (E) and a spline graph (F) of a mitochondrion with an isolation membrane elongating on its surface. The arrowhead indicates the elongating isolation membrane. The line graph represent the intensity profiles along the indicated lines in the respective images shown. Scale bars indicate 1 μm. See also Movie EV2 for the time-lapse video corresponding to (E).

Furthermore, when an isolation membrane started to engulf a mitochondrion, OPTN, which was initially distributed uniformly on the mitochondrial surface, underwent redistribution to the area contacting the isolation membrane (Fig. 4E, Movie EV2). OPTN signals on the isolation membrane-negative side of the mitochondrion diminished during this process (Fig. 4E and F). The OPTN-enriched region expanded together with the isolation membrane thereafter (Fig. 4E). These live-cell observations are consistent with the predictions of our mathematical models (Fig. 3). Taken together, our experimental data support the hypothesis that OPTN forms sheet-like phase-separated condensates on the surface of ubiquitinated mitochondria, exhibiting liquid-like properties.

### Liquid-like property of OPTN condensates is required for mitophagy

To investigate the importance of the liquid-like nature of OPTN condensates on mitochondria, we sought to reduce its liquidity by strengthening the interaction but reducing the multivalency between ubiquitin and GFP–OPTN constructs. To achieve this, we utilized the anti-green fluorescent protein (GFP) nanobody, which exhibits a high binding affinity to GFP (Rothbauer *et al*, 2008). Fusing the anti-GFP nanobody to the N-terminus of ubiquitin (nanobody–mRuby–Ub) (Fig. 5A, bottom panel) resulted in significantly enhanced binding to GFP–OPTN (Fig. 5B). Deleting the ubiquitin-binding domain in ABINs and NEMO (UBAN domain) from OPTN (GFP–OPTNΔUBAN) eliminated the interaction with ubiquitinated proteins (Fig. 5B). Wild-type GFP–OPTN accumulated on ubiquitinated mitochondria, but GFP–OPTNΔUBAN failed to do so (Fig. 5A, top and middle panels). In contrast, the expression of nanobody–mRuby–Ub rescued the recruitment of GFP–OPTNΔUBAN to mitochondria (Fig. 5A, lower panel). FRAP experiments revealed that, compared with wild-type GFP–OPTN (with mRuby–Ub, not nanobody-mRuby–Ub), GFP–OPTNΔUBAN showed a substantial decrease in recovery on mitochondria when co-expressed with nanobody–mRuby–Ub, indicating a reduction in its dynamic properties (Fig. 5C).

**Figure 5.**
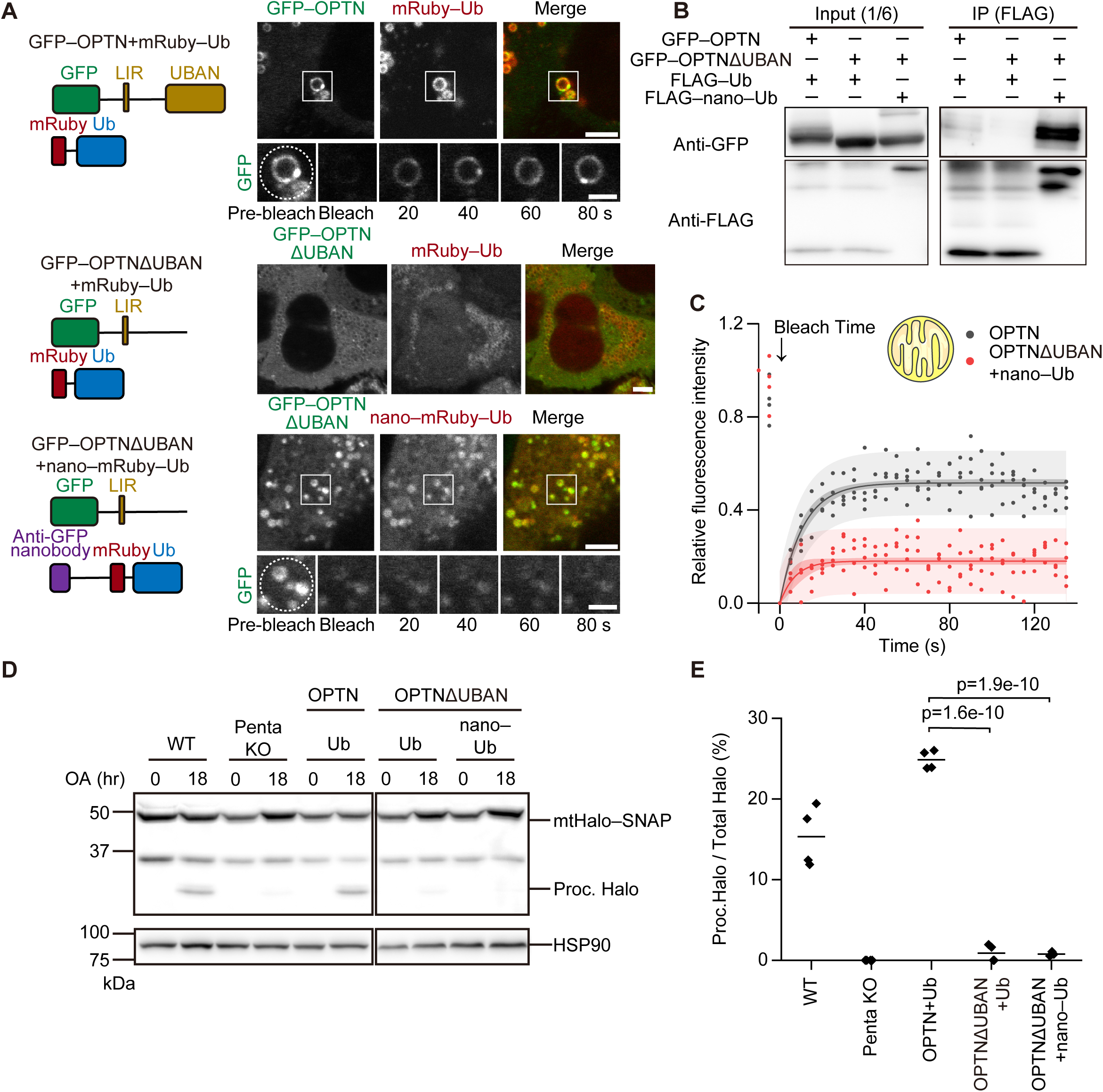
Liquid-like property of OPTN condensates is required for mitophagy. (A) HeLa cells lacking all five autophagy adaptors (penta KO cells) expressing both GPF– OPTN and mRuby–Ub (top), both GFP–OPTNΔUBAN and mRuby–Ub (middle), or both GFP–OPTNΔUBAN and anti-GFP nanobody–mRuby–Ub (bottom) at 45 min after 20 µM CCCP treatment. Time-lapse images of GFP FRAP are shown. Photobleached areas are circled by dotted lines. Scale bars indicate 4 μm and 2 μm (magnified images). (B) Interaction between GFP–OPTN mutants and FLAG–Ub or FLAG–nanobody–Ub was investigated by immunoprecipitation with an anti-FLAG antibody and immunoblotting with an anti-GFP antibody. (C) Quantification of GFP FRAP on separate mitochondria (Mt) in penta KO cells expressing both GFP–OPTN and mRuby–Ub or both GFP–OPTNΔUBAN and anti-GFP nanobody–mRuby–Ub at 45 min after 20 µM CCCP treatment. Data were collected from four structures and were fitted to the equation *y* = *a*\*(1 − exp(−*b*\**x*)). The dark shading represent the 95% confidence intervals, and the light shading represent the 95% prediction intervals. (D and E) Representative data (D) and quantification (E) of HaloTag (Halo) processing assay using cells expressing the indicated OPTN and Ub constructs. Cells expressing the mtHalo–SNAP mitophagy reporter were treated without (0 h) and with 1 μM oligomycin and 2 μM antimycin for 18 h. The amount of processed Halo (proc. Halo) indicates the relative amount of mitochondria degraded by the lysosomes. Solid horizontal bars indicate the means, and dots indicate the data from four independent experiments. Differences were statistically analyzed by one-way analysis of variance with Dunnett’s post-hoc test.

We then evaluated the effect of impaired OPTN liquidity on mitophagy using the recently developed HaloTag cleavage assay (Yim *et al*, 2022). This assay utilizes the ligand-dependent conformational change of HaloTag (Halo); ligand-free Halo is efficiently degraded in lysosomes, whereas ligand-bound Halo becomes resistant to lysosomal degradation. When Halo is expressed in the mitochondrial matrix by fusing to the mitochondrial presequence of Fo-ATPase subunit 9 and SNAP-tag (mtHalo–SNAP), we can measure the extent of mitochondrial degradation by quantifying the amount of processed free Halo out of the total amount of Halo (mtHalo–SNAP + processed Halo). Mitophagy activity, which was impaired in penta KO cells, was restored by the expression of GFP–OPTN (Fig. 5D and E). However, GFP–OPTNΔUBAN failed to restore mitophagy, even when it was co-expressed with nanobody–mRuby–Ub to rescue ubiquitin binding and mitochondrial localization (Fig. 5A, B, D, and E). These data suggest that the localization of OPTN on mitochondria and its binding to ubiquitin alone is insufficient for mitophagy, and the liquidity of OPTN is important for effective mitophagy.

### Liquid-like property of OPTN condensates is required for the recruitment of ATG9 vesicles

ATG9 vesicles are considered to be one of the sources of autophagosomal membranes (Yamamoto *et al*, 2012). Upon mitophagy induction, ATG9 vesicles accumulate at the sites of autophagosome formation (Itakura *et al*, 2012). OPTN plays a crucial role in this process by recruiting ATG9 vesicles to induce mitophagy (Yamano *et al*, 2020). In general, liquid-like condensates can be involved in clustering small vesicles such as synaptic vesicles, which exemplify complete wetting (Milovanovic & De Camilli, 2017; Sansevrino *et al*, 2023). Indeed, ATG9 vesicles are incorporated into condensates containing a glaucoma-associated OPTN mutant (O’Loughlin *et al*, 2020) and synapsin (Park *et al*, 2023). We, therefore, hypothesized that the liquid-like properties of OPTN condensates on mitochondria facilitate the recruitment of ATG9 vesicles. Consistent with previous reports (Itakura *et al*, 2012; Yamano *et al*, 2020), ATG9A was recruited to depolarized mitochondria in cells expressing GFP–OPTN (Fig. 6A and B). However, this ATG9A recruitment was almost completely abolished in cells expressing nanobody–mRuby–Ub and GFP– OPTNΔUBAN (Fig. 6A and B), even though GFP–OPTNΔUBAN still retains the ATG9A-interacting domain and the ability to bind with the ATG9 vesicles (Fig. 6C and D). Taken together, these findings suggest that the liquidity of sheet-like OPTN condensates on the mitochondrial surface is crucial for recruiting ATG9 vesicles and executing mitophagy.

**Figure 6.**
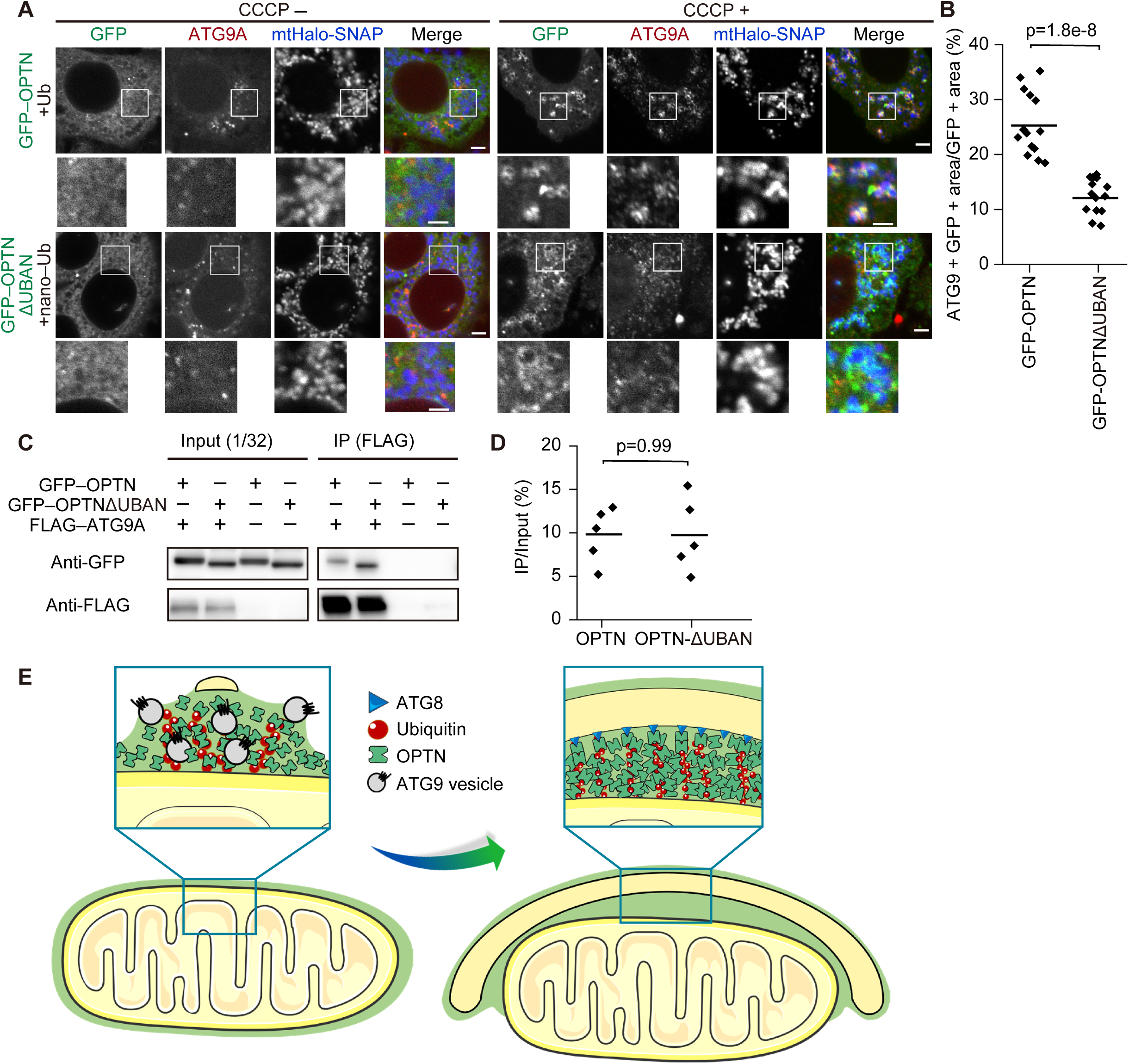
Liquid-like property of OPTN condensates is required for the recruitment of ATG9 vesicles. (A and B) Localization of ATG9 vesicles under normal (CCCP−) or mitophagy-inducing conditions (CCCP+, 60 min) (A) and quantification (B) of ATG9A colocalization with mitochondria. Endogenous ATG9 was immunostained in penta KO cells expressing both GFP–OPTN and mRuby–Ub or both GFP–OPTNΔUBAN and mRuby–nanobody–Ub together with mitochondrially targeted Halo–SNAP (mtHalo–SNAP). Scale bars indicate 4 μm and 2 μm (magnified images). Solid horizontal bars indicate the means, each dot indicates the mean value from one field of view with ≥10 cells (n=15). Differences were statistically analyzed by one-way analysis of variance with Dunnett’s post-hoc test. (C and D) Representative data (C) and quantification (D) of organelle immunoprecipitation in cells expressing GFP–OPTN or GFP–OPTNΔUBAN in the presence or absence of FLAG-ATG9A. ATG9 vesicles were immunoprecipitated by FLAG M2 beads and GFP was blotted to detect the interaction between GFP–OPTN (WT or ΔUBAN) and ATG9 vesicles at 60 min after 20 µM CCCP treatment. Solid horizontal bars indicate the means, and dots indicate the data from five independent experiments. Differences were statistically analyzed by one-way analysis of variance with Dunnett’s post-hoc test. (E) The models of ATG9 vesicle recruitment (left) and isolation membrane elongation (right) mediated by liquid-like OPTN condensates. Partial or complete wetting of phase-separated OPTN enables the recruitment of ATG9 vesicles to ubiquitinated mitochondria (left). Wetting of OPTN on mitochondria and isolation membranes also facilitates the engulfment of mitochondria by the isolation membranes (right).

## Discussion

### LLPS by autophagy adaptors during Parkin-mediated mitophagy

In the present study, we discovered that autophagy adaptors accumulate on ubiquitinated mitochondria with non-stoichiometric distribution compared with ubiquitin signals. They displayed characteristics consistent with phase-separated structures, including dynamic exchange with cytosol and redistribution upon coalescence. Notably, we observed the enrichment of OPTN at the isolation membrane-positive region, which expanded along with the elongation of the isolation membrane. These findings suggest that phase-separated adaptors facilitate the effective engulfment of mitochondria by the isolation membrane through the wetting effect and capillary forces (Fig. 6E, right panel). Moreover, the liquidity of OPTN condensates is critical for the localization of ATG9 vesicles onto ubiquitinated mitochondria, indicating that partial or complete wetting of phase-separated adaptors to ATG9 vesicles may be crucial for their recruitment to and/or retention on mitochondria (Fig. 6E, left panel). Therefore, we propose that LLPS of autophagy adaptors plays a critical role in two distinct steps of mitophagy by producing capillary forces, first, at the initiation step by recruiting and retaining the ATG9 vesicles in the condensates and, second, at the autophagosome elongation step by facilitating the attachment between ubiquitinated mitochondria and the isolation membrane.

### LLPS in bulk and selective autophagy

Recent reports have shed light on the importance of LLPS in selective autophagy (Noda *et al*, 2020). In lysophagy, p62 undergoes LLPS on lysosomes to facilitate efficient lysophagy together with HSP27, which maintains the liquidity of p62 and prevents its gelation (Gallagher & Holzbaur, 2023). This may also be mediated by the wetting between damaged lysosomes and the isolation membrane by phase-separated p62. Indeed, p62 itself forms phase-separated condensates along with ubiquitinated proteins, producing capillary forces by the wetting effect to facilitate the engulfment by the isolation membrane (Agudo-Canalejo *et al*, 2021). Similarly, Ape1, the cargo of the cytosol-to-vacuole targeting (Cvt) pathway in yeast, forms gel-like condensates with Atg19 on the surface of Ape1 condensates to be sequestered by the isolation membrane (Yamasaki *et al*, 2020). This association could also be mediated by partial wetting. These examples together with our observations highlight the importance of LLPS and the wetting phenomena in various selective autophagy processes.

LLPS may also be important for non-selective bulk autophagy. In yeast, the Atg1 complex (the ULK1 complex in mammals) forms pre-autophagosomal structures (PASs), which have been shown to be condensates driven by LLPS (Fujioka *et al*, 2020). Phase separation in this context provides a high local concentration of the Atg1 complex to support the autoactivation of Atg1 kinase (Yamamoto *et al*, 2016). Mammalian autophagy factors also accumulate at the autophagosome formation site; FIP200, a subunit of the autophagy-initiation complex, undergoes LLPS triggered by calcium transients (Zheng *et al*, 2022). The liquid-like nature of these autophagy-initiating structures may also be important for the recruitment of Atg9/ATG9 vesicles through the wetting effect.

### The importance of phase separation during mitophagy

In contrast to Parkin-mediated mitophagy, hypoxia-induced mitophagy does not involve autophagy adaptors, and instead, the outer mitochondrial membrane proteins NIX (also known as BNIP3L) and BNIP3 directly interact with ATG8 homologs on the isolation membrane (Novak *et al*, 2010; Onishi *et al*, 2021). In this context, ATG8 proteins interact with NIX or BNIP3, likely in a one-to-one manner. This raises the problem of why phase separation of autophagy adaptors is essential for Parkin-mediated mitophagy. One possibility is that the quick induction and execution of Parkin-mediated mitophagy requires amplification of the reactions through increased local concentration of adaptors and autophagy-related proteins facilitated by LLPS. This is consistent with the fact that mitophagy-inducing signals are also amplified by a mechanism involving phosphorylated ubiquitin (Kane *et al*, 2014; Kazlauskaite *et al*, 2014; Koyano *et al*, 2014). Alternatively, Parkin-mediated mitophagy requires a high level of precision and selectivity; mitophagy should be induced only for damaged mitochondria, which is ensured by the recruitment and retention of ATG9 vesicles only on phase-separated adaptor-positive mitochondria. This strict specificity would enable proper quality control of the mitochondria. Further study would be required to elucidate the functions and physiological importance of LLPS by autophagy adaptors in mitophagy and beyond.

## Materials and Methods

### Cell culture and generation of stable cell lines

HeLa cells, HeLa penta-knockout (penta KO) cells, and human embryonic kidney (HEK) 293T cells authenticated by RIKEN were cultured in Dulbecco’s modified Eagle’s medium (DMEM; D6546; Sigma-Aldrich) supplemented with 10% fetal bovine serum (FBS; 173012; Sigma-Aldrich) and 2 mM L-glutamine (25030-081; Gibco) in a 5% CO_2_ incubator at 37°C. For stable expression, retrovirus was produced with HEK293T cells transfected with pMRX-IP-based, pMRX-IB-based, pMRX-IU-based, or pMRX-IN-based retroviral plasmids, pCG-VSV-G, and pCG-gag-pol by using Lipofectamine 2000 (11668019; Thermo Fisher Scientific). After transfection, cells were further incubated at 37°C for 24h. The viral supernatant was collected by filtration through a 0.45-μm filter unit (Ultrafree-MC; Millipore) and then used for infection. Cells were plated onto 6-cm dishes 18 h before infection, and the medium was replaced with viral supernatant diluted 1.5-fold with 8 μg/mL polybrene (H9268; Sigma-Aldrich). Two days later, cells were selected in a medium containing 2 μg/mL puromycin (P8833; Sigma-Aldrich), 4 μg/mL blasticidin S hydrochloride (022-18713; FUJIFILM Wako Pure Chemical Corporation), 1.5 mg/mL neomycin, or 250 μg/mL zeocin (R25005; Thermo Fisher Scientific).

### Plasmids

The pMRX-IPU, pMRX-IBU and pMRX-INU plasmids were generated by modifying the multi-cloning site of pMRX-IP (Saitoh *et al*, 2002; Kitamura *et al*, 2003), pMRX-IB, and pMRX-IN, respectively. DNA fragments encoding ubiquitin, p62 (NM_003900.5), OPTN (NM_001008211.1), NBR1 (NM_001291571.2), NDP52 (NM_001261391.2), TAX1BP1 (NM_001206901.1), ATG9A (NM_001077198.3), LC3A (NM_032514.4), and LC3B (NM_022818.5) were inserted into pMRX-IPU, pMRX-INU, or pMRX-IBU. DNAs encoding the HA epitope, monomeric enhanced GFP with the A206K mutation (mGFP), codon-optimized ultra-stable GFP (muGFP) (Scott *et al*, 2018), codon-optimized mRuby3 (Bajar *et al*, 2016), HaloTag7 (G1891; Promega), 3×FLAG, and SNAP-tag (New England BioLabs, N9181S) were used for tagging. The mitochondrial presequence of *Neurospora crassa* Fo-ATPase subunit 9 (residues 1–69) was added to HaloTag7-SNAP to deliver the reporter into the mitochondrial matrix (mtHalo-SNAP) (Eura, 2003). Truncated OPTN (OPTNΔUBAN, aa 445–502) was prepared by PCR-mediated site-directed mutagenesis. The resulting plasmids were sequenced.

### Antibodies and reagents

The primary antibodies used in this study were rabbit polyclonal anti-OPTN (Proteintech, 10837-AP), rabbit polyclonal anti-ATG9A (MBL, PD042), mouse monoclonal anti-Halo (Promega, G9211), mouse monoclonal anti-HSP90 (BD Transduction Laboratories, 610419), and rabbit polyclonal anti-GFP (Thermo Fisher Scientific, A6455) antibodies. The secondary antibodies used were HRP-conjugated goat polyclonal anti-rabbit IgG (Jackson ImmunoResearch Laboratories, 111-035-144) and HRP-conjugated goat polyclonal anti-mouse IgG (Jackson ImmunoResearch Laboratories, 115-035-003) antibodies. To induce mitophagy, HeLa cells were treated with 10 μM carbonyl cyanide *m-*chlorophenyl hydrazine (CCCP; Sigma-Aldrich) for 45 min, or 10 μM oligomycin (Cabiochem, 495455-10MGCN) and 4 μM antimycin A (Sigma-Aldrich, A8674) for 18 h. After cells had been treated with oligomycin and antimycin A for >6 h, 10 μM Q-VD-OPH (SM Biochemicals, SMPH001) was added to block apoptotic cell death. To inhibit the formation of isolation membranes in the 1,6-hexanediol (Sigma-Aldrich, 240117-50G) treatment experiment, 20 μM wortmannin (Sigma-Aldrich, W1628-1MG) was added into the medium.

### Immunoprecipitation and immunoblotting

Cell lysates were prepared in a lysis buffer (50 mM Tris-HCl pH 7.4, 150 mM NaCl, 1 mM EDTA, 1% Triton X-100, EDTA-free protease inhibitor cocktail [19543200; Roche]). After centrifugation at 17,700 × *g* for 10 min, the supernatants were subjected to immunoprecipitation using anti-FLAG M2 magnetic beads (M8823-1ML; Sigma-Aldrich). Precipitated immunocomplexes were washed three times with lysis buffer and boiled in sample buffer (46.7 mM Tris-HCl, pH 6.8, 5% glycerol, 1.67% sodium dodecyl sulfate, 1.55% dithiothreitol, and 0.02% bromophenol blue). For immunoprecipitation of ATG9 vesicles, cells were disrupted by Dounce homogenization with hypotonic lysis buffer (10 mM HEPES, pH 7.9, 1.5 mM MgCl_2_ and 10 mM KCl, EDTA-free protease inhibitor cocktail [19543200; Roche]). Disruption was carried out by applying 35 strokes while the cell suspension was cooled on ice. The supernatants were subjected to immunoprecipitation using anti-FLAG M2 magnetic beads (M8823-1ML; Sigma-Aldrich). For immunoblotting, the samples were separated by SDS-PAGE and transferred to Immobilon-P polyvinylidene difluoride membranes (Millipore, WBKLS0500) with the Trans-Blot Turbo Transfer System (Bio-Rad). After incubation with the relevant antibody in 5% skim milk in 20 mM Tris-HCl, 150 mM NaCl, and 0.1% Tween 20 (02194841-CF; MP Biomedicals), the signals from incubation with SuperSignal West Pico Chemiluminescent Substrate (Thermo Fisher Scientific, 34579) were detected with the FUSION SOLO.7S.EDGE imaging system (Vilber-Lourmat). Contrast and brightness adjustment and quantification were performed using the image processing software Fiji (Schindelin *et al*, 2012).

### Fluorescence recovery after photobleaching

In-cell fluorescence recovery after photobleaching (FRAP) analyses were performed with an Olympus Fluoview FV3000 confocal microscope equipped with a 60× oil-immersion objective lens (1.40 NA, Olympus). The chamber was maintained at 37°C and continuously supplied with humidified 5% CO_2_. Bleaching was performed using 80% laser power (488 or 561 nm laser), and images were captured every 5 s for 30 frames. Recovery curves and fitting were analyzed using OriginPro 2022. Fluorescence intensity of the bleached spot, an unbleached control spot, and the background was measured using the software package Fiji. Background intensity was subtracted, and the intensity values of the region of interest are reported relative to the pre-bleached images during image acquisition. Each data point represents the mean and standard error of fluorescence intensities in more than three unbleached (control) or bleached (experimental) spots.

### Live-cell imaging

Living imaging was conducted with the Olympus SpinSR10 spinning-disk confocal super-resolution microscope equipped with a Hamamatsu ORCA-Flash 4.0 camera, a UPLAPO OHR 100 × (NA 1.50) lens, and the SORA disk in place. The microscope was operated with Olympus cellSens Dimension 2.3 software. Cells were passaged onto a four-chamber glass-bottom dish (Greiner Bio-One) more than 24 h before imaging. To induce mitophagy, cells were incubated with 10 μM CCCP in the presence of 200 nM SF650-conjugated Halo ligand (GoryoChemical, A308-02). Images were processed using the image processing software package Fiji (Schindelin *et al*, 2012). Fluorescence intensity of indicated fluorophores was measured by Fiji, and spline connected graphs were created using OriginPro 2022 software.

### Immunofluorescence imaging

Cells were fixed with 4% PFA in 0.1 M phosphate buffer (pH 7.3) for 15 min at room temperature (RT) and washed with PBS. Fixed cells were incubated with 10 µg/mL digitonin for 5 min at RT and washed with PBS. Then, cells were incubated with primary antibodies diluted (1:1000) in blocking buffer (3% BSA, in PBS) for 1 h at RT, washed with PBS, incubated with Alexa Fluor-conjugated secondary antibodies (1:1000) in blocking buffer for 60 min at RT, washed again, and mounted on coverslips. Colocalization was calculated by Fiji plugins BIOP JACoP, with the threshold of the ATG9A channel fixed to 500. ATG9A-positive areas out of GFP-positive areas were calculated to obtain the overlapping areas.

### Mathematical model of autophagy adaptor condensate formation

We formulated a mathematical model of autophagy adaptor condensate formation. For the case of two mitochondria approaching each other (Fig. 3A), the free energy, *F*, of the system can be written as

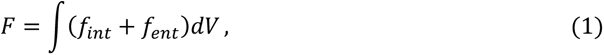

with the interaction energy, *f_int_*, and the entropic energy, *f_ent_*, expressed as

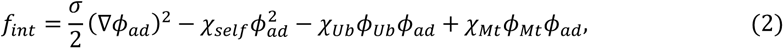

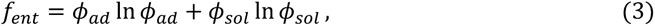

where *ϕ_ad_*, *ϕ_Ub_*, and *ϕ_sol_* = 1 − *ϕ_ad_* − *ϕ_Ub_* − *ϕ_Mt_* are the adaptor, ubiquitin, and solvent concentrations, respectively, *χ_self_*and *χ_Ub_* represent the strength of the adaptor self-interaction and adaptor–ubiquitin interaction, respectively, σ represents the surface tension of the condensates, and *χ_Mt_* represents the mitochondrial area exclusion effect, introduced to prevent proteins from entering the mitochondrial interior (*ϕ_Mt_*).

For the case in which an isolation membrane surrounded a mitochondrion (Fig. 3C), the interaction energy, *f_int_*, is instead expressed as

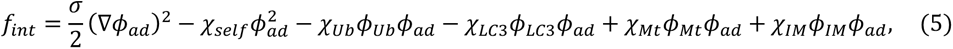

where *ϕ_LC3_* and *χ_LC3_* represent the LC3 concentrations and the strength of the adaptor–LC3 interaction, respectively, and *χ_IM_* represents the mitochondrial area exclusion effect, introduced to prevent proteins from entering the lumen of the isolation membrane (*ϕ_IM_*). The solvent concentration that appears in the entropic energy (3) was also modified to

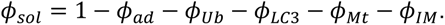

For the numerical simulation, the space was discretized by a lattice with 10 nm on each side, and a concentration field was assigned to each location. The time evolution of the system was assumed to follow the Cahn–Hilliard equation,

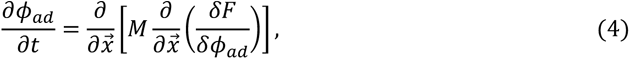

which is commonly used in analyses of phase separation dynamics (Berry *et al*, 2018). Here, *M* = *D*/*k_B_T* is the mobility of the protein with the diffusion constant *D* = 10 μm^2^/sec (Milo & Phillips, 2015) expressed in terms of the Boltzmann constant *k_B_* and temperature *T*. We considered a 3000 nm × 3000 nm square region with periodic boundary conditions containing two mitochondria and surrounding cytosolic components as a system and assumed that adaptor proteins are uniformly distributed in the cytosol around mitochondria at a concentration of *ϕ* = 0.1 in the initial state.

### Statistical analysis

Statistical analysis was performed using OriginPro 2022 software. The statistical methods used for each analysis are specified in the figure legends.

## Acknowledgments

We thank Michael Lazarou (Walter and Eliza Hall Institute of Medical Research) for providing pentaKO HeLa cells, Toshio Kitamura for pMXs-IP (The University of Tokyo), Shoji Yamaoka (Tokyo Medical and Dental University, Tokyo, Japan) for pMRXIP, and Teruhito Yasui (National Institutes of Biomedical Innovation, Health and Nutrition (NIBIOHN), Osaka, Japan) for pCG-VSV-G and pCG-gag-pol. pKanCMV-mClover3-mRuby3 was a gift from Michael Lin (Addgene plasmid # 74252 ; http://n2t.net/addgene:74252 ; RRID:Addgene_74252).

This work was supported by the Exploratory Research for Advanced Technology (ERATO) research funding program of the Japan Science and Technology Agency (JST) (JPMJER1702 to NM), a Grant-in-Aid for Specially Promoted Research (22H04919 to NM) and a Grant-in-Aid for Scientific Research (C) (23K05715 to Y.S.) from the Japan Society for the Promotion of Science (JSPS), and German Research Foundation (Project 460056461 to RLK). The numerical computations have been performed with the RIKEN supercomputer system (HOKUSAI).

## Author contributions

Z.Y., S.Y., H.C. R.L.K., and N.M. designed the project. Z.Y. performed most of the cell biology experiments. Y.S. developed the mathematical model. Z.Y., S.Y., Y.S., R.L.K., and N.M. wrote the manuscript. All authors analyzed and discussed the results and commented on the manuscript.

**Movie EV1. OPTN condensates redistribute upon mitochondrial contact.**

Time-lapse video of two separate mitochondria approaching and contacting each other (30s per frame). Scale bars indicate 1 μm. See also Figure 4C and D.

**Movie EV2. OPTN condensates redistribute upon mitochondria–isolation membrane contact.**

Time-lapse video of a mitochondrion with an isolation membrane elongating on its surface (30s per frame). Scale bars indicate 1 μm. See also Figure 4E and F.

---

**Figure EV1.**
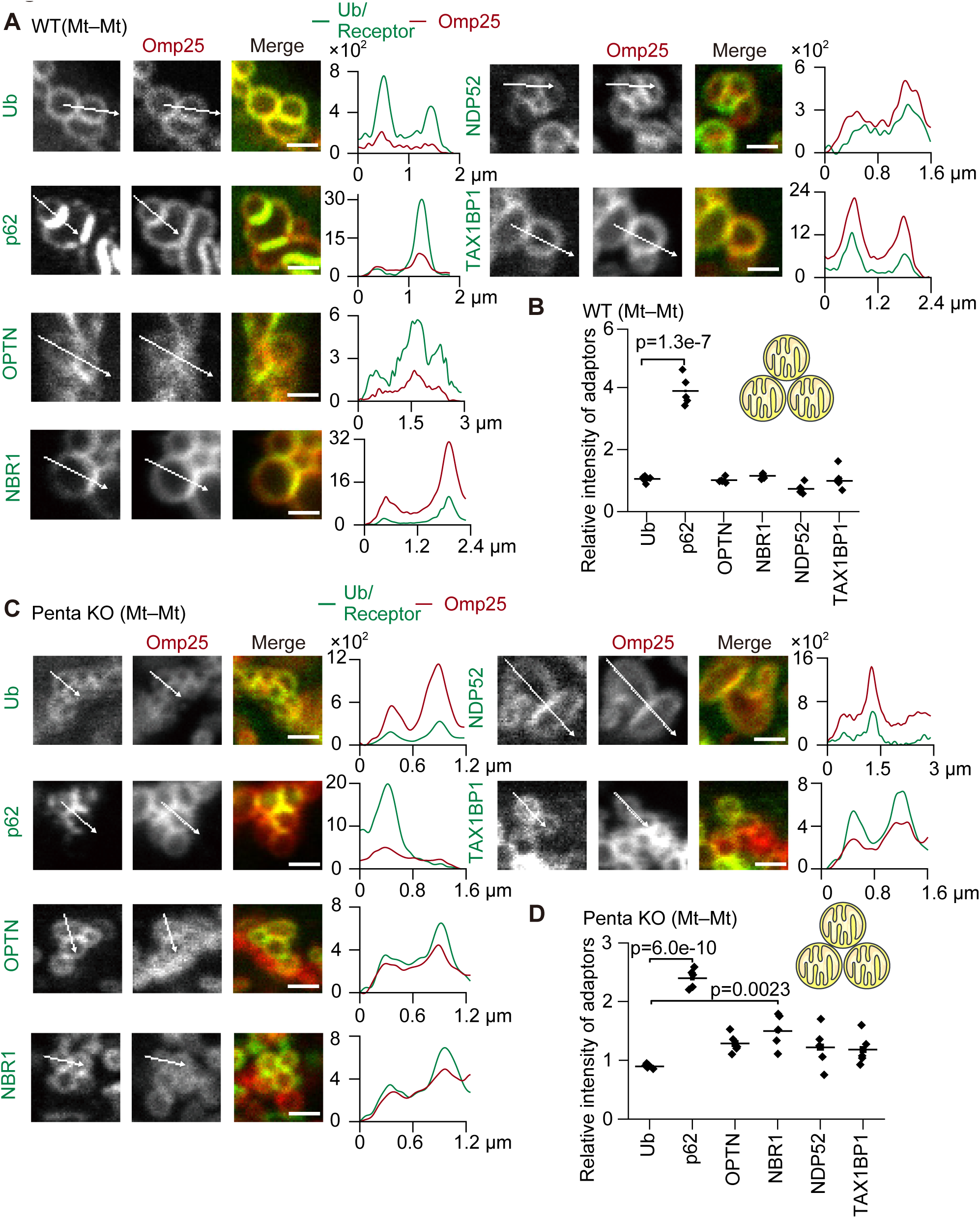
Localization of autophagy adaptors between mitochondria during Parkin-mediated mitophagy. (A and C) Representative images (left) and spline graphs of the intensity profiles along the indicated lines (right) of wild-type HeLa cells (A) or HeLa cells lacking all five autophagy adaptors (penta KO cells) (C) expressing one of the GFP–tagged autophagy adaptors or ubiquitin and mRuby–Omp25 at 45 min after 20 µM CCCP treatment. Mitochondrial clusters (Mt–Mt) are shown. The *y*-axis in each of the graphs indicates the fluorescence intensity. (B and D) The relative intensity of each adaptor was calculated as in Figure 1C. Solid horizontal bars indicate the means, and dots indicate the data from five structures. Differences were statistically analyzed by one-way ANOVA with Dunnett’s post-hoc test.

